# Enhanced expression of 4-Cl-IAA and 6-Cl-IAA by touch stimulus for rapid and differential growths of the Madeira vine

**DOI:** 10.1101/2022.04.13.488269

**Authors:** Ma-Hsuan Ma, Erdembayalag Batsaikhan, Chun-Ming Wu, Hao-Hsun Lee, Chih-I Luo, Ni-Jhen Chen, Jeng-Der Chung, Ching-Te Chien, Yu-Han Tsai, Wen-Hsien Li

**Author notes:** Authors Ma-Hsuan Ma, Erdembayalag Batsaikhan, Chun-Min Wu, Hao-Hsun Lee, Chih-I Luo, Ni-Jhen Chen, Jeng-Der Chung Ching-Te Chien, Yu-Han Tsai.

## Abstract

Madeira vine (MV) grows 30 times faster after encountering a support. In vivo x-ray diffraction made on live MV stems revel the appearance of crystallized IAA (C_10_H_9_NO_2_), 4-Cl-IAA (C_10_H_8_ClNO_2_) and 6-Cl-IAA (C_10_H_8_ClNO_2_) in the stems. Small angle neutron scattering spectra of the IAA extracted from MVs stem reveal a progressive increase in the size of crystallized IAA transported downward from the apex of the shoots. High resolution X-ray diffractions made on the extracted IAA reveal significantly larger amounts of 4-Cl-IAA and 6-Cl-IAA in the climbing MVs than in the swaying around MVs. The gas chromatography-mass spectrometry spectra reveal the production of 9% more IAA and 90% more 4-Cl-IAA+6-Cl-IAA at the apexes of climbing MVs than swaying MVs. More 4-Cl-IAA+6-Cl-IAA were transported to the contact-free side than to the contact side of the vine. In vivo neutron tomography of naturally climbing MVs reveals a substantially higher H^+^ concentration in the contact-free parts than in the contact parts. The absorption spectra also reveal more expansin in the contact-free parts than in the contact parts. These results provide a view, at the molecular level, of what triggers the faster and differential growths in MVs in response to touching a support.

## Introduction

Although plants are unable to move as quickly as animals, they are very much in tune with their environment and are capable of a variety of movements (Steinitz and Hagiladi, 1987; Braam, 2005; Esmon et al., 2005; Rowe and Speck, 2005; Iino, 2006; Bowling and Vaughn, 2009). Among them, swaying followed by climbing or twining around a support is characteristic of the growth of vine plants (Guerra et al., 2019). The tip of a growing vine stem actually swings around in wide circles, a process known as circumnutation (Stolarz, 2009; Smyth, 2016), until it encounters a support to wind around as it grows. After twining around a support, vines invest less energy in their own supportive tissue, but spend more on growing faster towards the sunlight (Gianoli, 2015; Burris et al., 2018; Zhang et al., 2018). This behavior which occurs in response to mechanical contact is known as thigmotropism, derived from the Greek words thigmo for touch and tropism for turning. Thigmotropism was first described by Charles Darwin in his book “*The power of movement in plants*” published in 1880 (Darwin, 1881). The thigmotropic response triggers two actions, faster growth and differential growth in the growing stem (Darwin, 1881; Jaffe et al., 2002). Faster stem growth is a direct result of intense enlargement of the cells in the stem caused by transport of more auxins into the cells. Differential growth is marked by the production of longer and stronger cells on the contact-free (CF) parts than on the contact (CT) parts of the stem as it winds around the support. This process requires an adjustment of the growth chemistry and biomass reallocation across the stem to provide a better growth environment for the CF parts than for the CT parts.

For a long time after plant tropisms were discussed by Darwin in 1880, little was known about thigmotropism due to the technical difficulty of separately measuring the physiological conditions in the CF and CT parts of a curled stem. It is now known is that it is the accumulation of the plant hormone, auxin indole-3-acetic acid (IAA) C_10_H_9_NO_2_, in the CF parts of the stem which drives the CF parts to grow faster than the CT parts, encouraging coiling onto its stimulator (Jaffe and Galston, 1968; Bopp and Weber, 1981; Lee et al., 2020). However, the question as to “how the vine can provide sufficient amounts of active IAA and effectively transport it into the cells for fast growth in such a short time” has not yet been answered. In response to this question, using Anredera cordifolia, also known as Madeira vine (MV), as an example, we look into the biological actions triggered by physical touch at the molecular level, using scattering and imaging techniques for in vivo and in vitro investigations. Anredera cordifolia (Fig. 1), is an evergreen vine originating from South America but now distributed worldwide, belonging to the Basellaceae family. MV is a fast climber, growing from fleshy rhizomes. Day-to-day measurements reveal that the daily growth rate can be as much as 18 times faster after a support is encountered, while the nighttime growth rate can be 4 times faster than that in daytime. In vivo x-ray diffraction (XRD) measurements reveal the appearance of crystallized IAA (C_10_H_9_NO_2_) and 4-Cl-IAA (C_10_H_8_ClNO_2_) in the MV stems. Mass spectra of auxin extracted from the stems reveal the generation of 20% more auxin by the apex in the climbing stems. Touch stimulus greatly enhances the expression of the highly active 4-Cl-IAA and 6-Cl-IAA (C_10_H_8_ClNO_2_) stimulating faster growth. The in vivo neutron images reveal a substantially higher H concentration in the CF parts than in the CT parts. The absorption spectra of crushed fresh MV stems reveal more expansin in the CF parts than in the CT parts. These results reveal the biological response of MVs to encountering a support for a rapid growth.

**Fig. 1.**
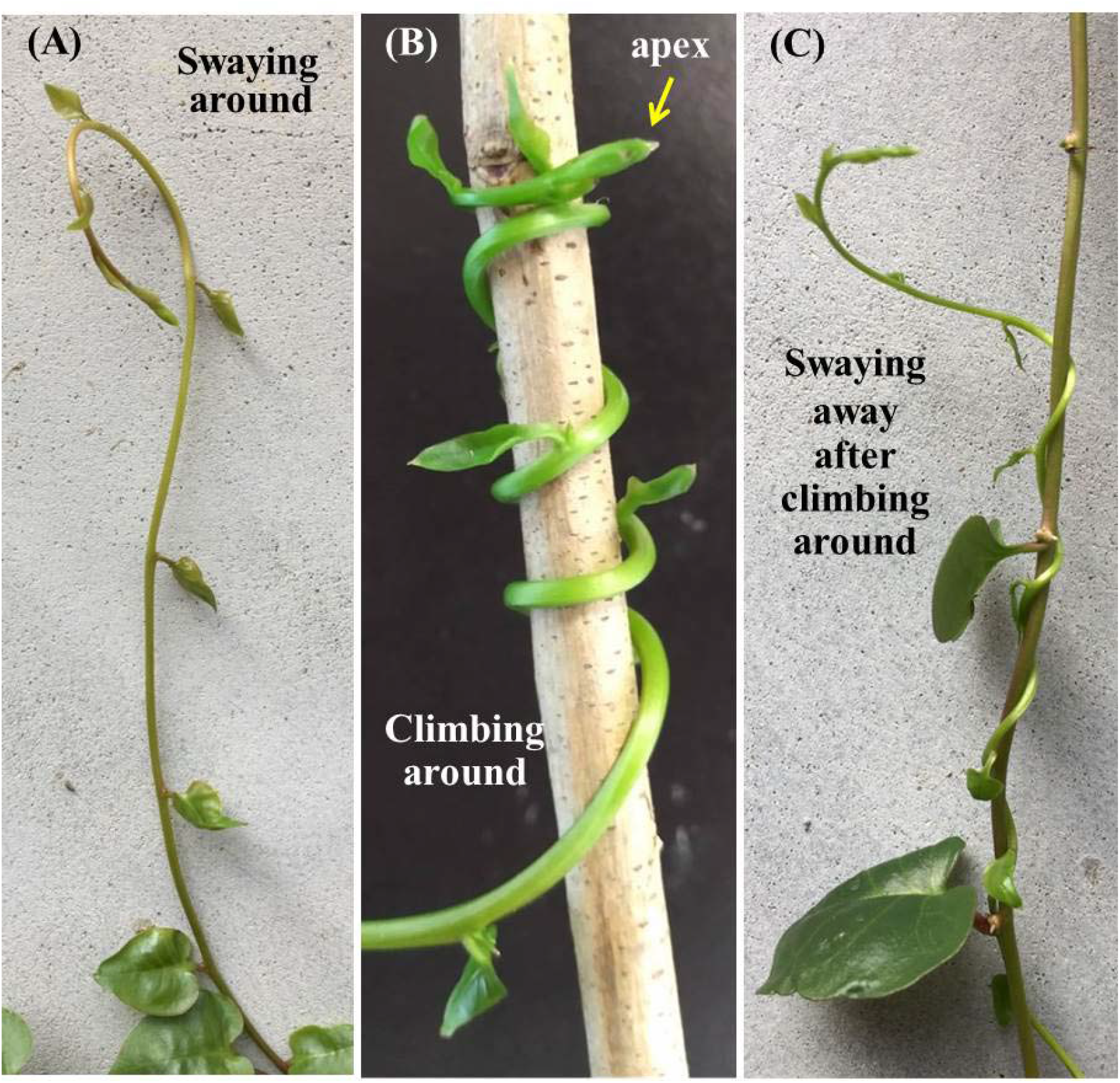
Growth patterns of Madeira vines. (A) A MV swaying around counterclockwise searching for a support. (B) A MV twining around a support after encountering it. (C) A MV swaying away from the support after twining around it for several days.

## Materials and methods

### X-ray diffraction

The X-ray diffraction measurements for structural analysis were performed on a Bruker D8 ADVANCE diffractometer. The in vivo diffraction on live MVs was performed using transmission geometry, with the beam incident angle fixed at zero throughout the scans. Each diffraction measurement took 3 hours to scan a 3 degree scattering angle, followed by a 6 hour beam off period prior to taking the next 3 hour diffraction measurement. The diffraction made on the extracted powders were performed employing the standard reflection geometry, where the beam incident angle and the reflective angle were set to be equal throughout the scans.

### IAA powder extraction

The IAAs in the swaying MVs were extracted from 150 stems collected in the field. Each stem was cut from the shoot apex into 1 cm long sections. Five groups of severed stems were collected: 0-1 cm (marked SA01), 2-3 cm (marked SA23), 4-5 cm (marked SA45), 6-7 cm (marked SA67), and 8-9 cm (marked SA89) from the apex. The IAAs in the climbing MVs were extracted from 95 stems, collected in the field. Three specific portions of each stem were sectioned: 0-1 cm from the shoot apex in the straight (ST) part of the stems; 0-5 cm from the contact point in the curved stems bisected into contact (CT) and contact-free (CF) sides. Extraction was performed using the High Performance Liquid Chromatography (HPLC) system, manufactured by Beckman. A commercial IAA powder (from Sigma-Aldrich) was used to mark the time channels for IAA collection. The extraction was carried out after mixing the crushed (at liquid N_2_ temperature) and dried (in purified N_2_ gas flow) stems (~0.3 g each) into a solution containing 15 ml of 80% (v/v) methanol, 0.4 mg butylated hydroxytoluene and 2 mg ascorbate at 4 °C for 15 hrs. The samples thus extracted were in powdered forms.

### Gas chromatography–mass spectrometry (GC-MS)

GC-MS for mass analysis was performed on an Agilent HP6890 Gas Chromatograph, equipped a 5793 MSD mass detector, with the sample loaded on a DB-1 capillary column (30 m in length × 0.25 mm inner diameter, 0.25 μm film thickness) from J&W Scientific. The time channel in the GC-MS spectrum for IAA was identified using a commercial IAA powder (from Sigma-Aldrich).

### Small angle neutron scattering (SANS)

The SANS measurements were performed on QUOKKA at the Bragg Institute, ANSTO, employing a velocity selector to extract neutrons of wavelength λ = 5 Å, a source-to-sample distance of 8 m, an evacuated flight path of 8 m between the sample and the detector, and a ^3^He two-dimensional position sensitive detector. For SANS measurements, the extracted powders (~10 mg each) were dispersed in D_2_O and loaded into a quartz cell mounted in a liquid-circulating temperature controller, operated at 20 or 40 °C. The intensities of the scattered neutrons from the sample and from the D_2_O buffer in quartz cell were measured. The scattering intensities from the IAA/4-Cl-IAA were isolated from the background by subtracting the buffer pattern from the sample pattern. The resulting two-dimensional data arrays were then radically averaged to obtain the intensity profile *I*(Q), where Q is the wavevector transfer.

### Neutron Tomography

The neutron tomographic images were taken on DINGO at the Bragg Institute, ANSTO, employing the high-intensity mode with a ZnS/LiF scintillator to achieve a field-of-view of 115×197 mm with 39×39 μm pixels.

## Results

### Faster growth

In this study, faster growth induced by mechanical touch was assessed by measuring the length from the tip of the shoot to a mark made at 1 cm below the tip at time zero. Each of ten MVs growing wild in a field, near Xiangshan in north Taiwan, measured twice daily at 7 am and 7 pm for 11 consecutive days. Among them, four MVs were swaying around (SA) without touching any support (namely the SA stems), three were climbing around (CA) a support but began swaying away from the support after several days (namely the CA stems), and the other three did not match the above categories. The SA stems were found to have a growth rate of 2.4 mm/day (open squares in Fig. 2A). Faster growth could be seen in the CA stems. Measurements made at 7 am (filled circles in Fig. 2) revealed more growth than those made at 7 pm (open triangles in Fig. 2), showing growth during the nighttime (7 pm to 7 am) to be much faster than growth during the daytime (7 am to 7 pm). The daytime growth rate remained at ~9 mm in 12 hrs during the climbing process (filled circles in Figs. 2B, 2C, 2D), which was still twice as fast as that of the SA stems. Once the vine touched a support, the nighttime growth increased daily, reaching a saturation maximum of ~35 mm in 12 hrs after climbing on a support for 3 days, but reducing to a rate essentially the same as the daytime growth rate 1 day after the apex swayed away from the support (open triangles in Figs. 2B, 2C, 2D). Interestingly, the faster growth continued at a reduced rate for ~1 day after the apex swayed away from the support, which amounted to an additional growth of ~20 mm. Apparently, touch induced climbing of MV triggers a growth rate that can be 35/1.2 = 30 times faster in the nighttime and 9/1.2 = 7.5 times faster in the daytime, and it took an ~3 day period for the plant to adjust to the growth environment in response to mechanical touch but only ~1 day to sense the loss of touch.

**Fig. 2.**
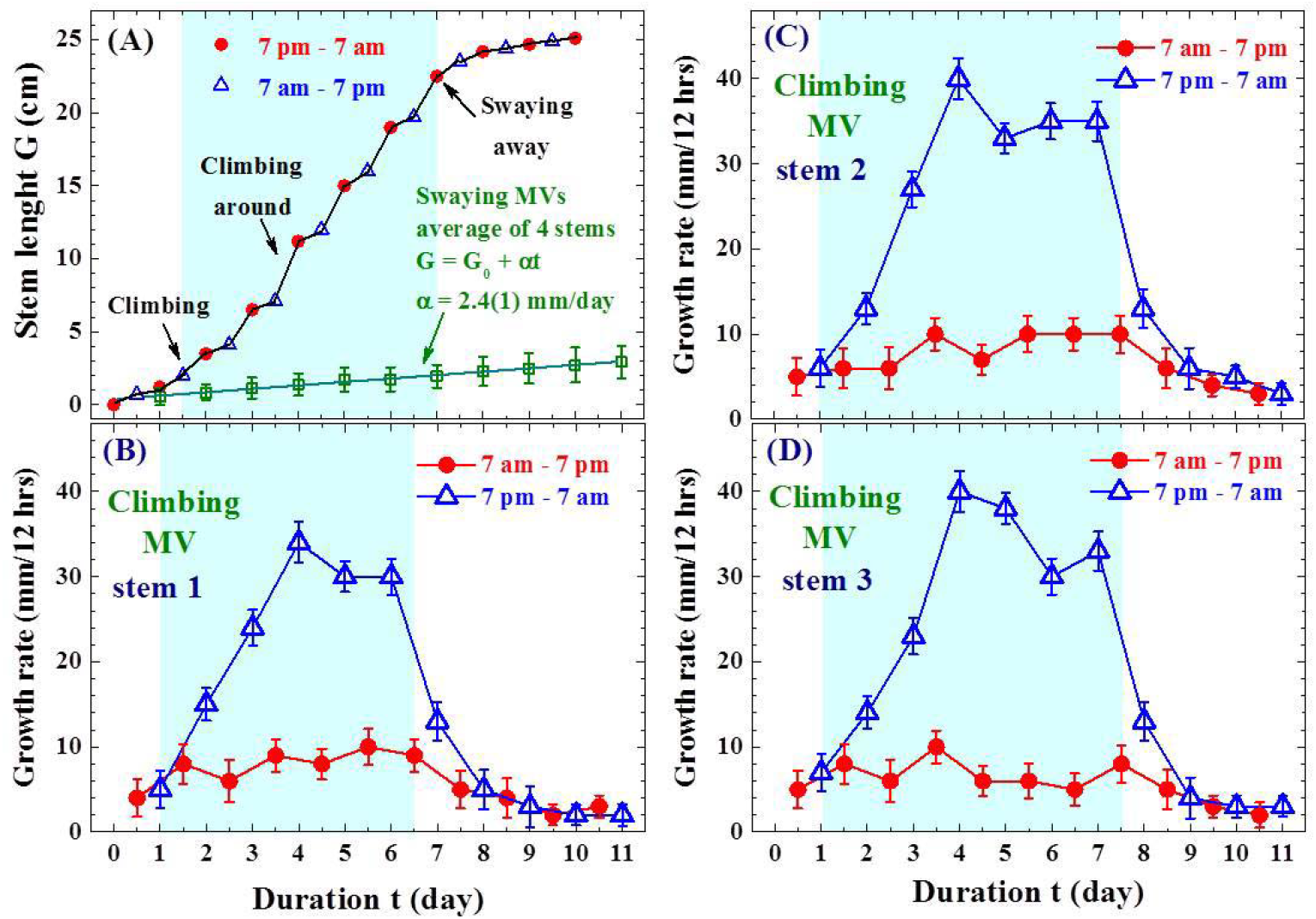
Daytime and nighttime growth rates. Measurements made on the length of a stem from the shoot tip to a mark placed at time zero. Blue shaded regions indicate the durations for which stems were twining around the supports. (A) Comparison between growth rate of a climbing (filled circles and open triangles) and swaying (open filled squares) stems, revealing much faster growth of the twining stem. (B)-(D) Daytime (solid circles) and nighttime (open triangles) growth rates of three separate twining stems, revealing much faster growth in the nighttime (7 pm to 7 am) than in the daytime (7 am to 7 pm).

### Structural form of active auxin – revealed by in vivo x-ray diffraction

Plant growth and morphogenesis are driven by the plant hormone auxin. Three naturally occurring active auxins, indole-3-acetic acid (IAA, C_10_H_9_NO_2_), 4-chloroindole-3-acetic acid (4-Cl-IAA, C_10_H_8_ClNO_2_), and phenylacetic acid (PAA, C_8_H_8_O_2_), have been identified in plant (Korasick et al., 2013). Among them, IAA is the most frequently found active auxin and has been recognized to be the key ingredient for cell enlargement and division in plants (Perrot-Rechenmann, 2010; Scorza and Dornelas, 2015), participating directly in all tropism responses (Scorza and Dornelas, 2015; Živanović et al., 2018). An IAA molecule consists of two benzenes and a carboxyl group, where the carboxyl group is weakly connected to the benzene group. IAA loses its ability to stimulate cell growth once the carboxyl group is detached, thus becoming a part of the inactive auxin pool (Korasick et al., 2013). 4-Cl-IAA is a chlorinated analogue of IAA. IAA diverges to 4-Cl-IAA when the amino acid tryptophan is chlorinated to form 4-chlorotryptophan (Živanović et al., 2018; Tivendale et al., 2012). Auxin is produced in the growing shoot tips and transported downward to the main stem via vascular bundles (Bennett et al., 2016). Once produced, it is distributed throughout the plant via a cell-to-cell transport system through localized plasma membrane transporter proteins (Vieten et al., 2007; Gälweiler et al., 1998; Bennett et al., 2006). Although it is believed that IAAs take their molecular form while being transported through the stem and cells, no investigation to prove or disapprove this general belief has yet been carried out.

Direct in vivo x-ray diffraction (XRD) measurements were made on the 0-1 cm from the shoot tips of a bundle of four live SA (Fig. 3A) and CA (Fig. 3E) MVs. The XRD pattern reveal well-defined Bragg diffraction peaks (Figs. 3B-3D and 3F). Well-defined XRD peaks belonging to the (100), (200) and (300) Bragg reflections of the IAA structure as well as the (100) and (120) Bragg reflections of the 4-Cl-IAA structure can be identified in the diffraction patterns taken on the SA stems (Figs. 3B-3D). The appearance of these well-defined diffraction peaks shows that the IAA and 4-Cl-IAA molecules in the MVs are linked into a periodic structure, rather than each remaining in molecular form. Three additional diffraction peaks (arrows in Fig. 3F), identified as the (210), (020) and (220) Bragg reflections of the 6-Cl-IAA structure (Nigović et al., 1996), appear in the diffraction patterns taken on the CA stems. The diffraction intensities from this component are weak but still evident in the diffraction patterns of the SA stem (Fig. 3B). Apparently, there is more 6-Cl-IAA in the CA stems than in the SA stems. Note that it has been demonstrated that both 4-Cl-IAA and 6-Cl-IAA are much more active than IAA in stimulating the growth reaction (Nigović et al., 1996; Böttger et al., 1978; Karcz and Burdach, 2002; Pless et al., 1984; Ahmad et al., 1987).

**Fig. 3.**
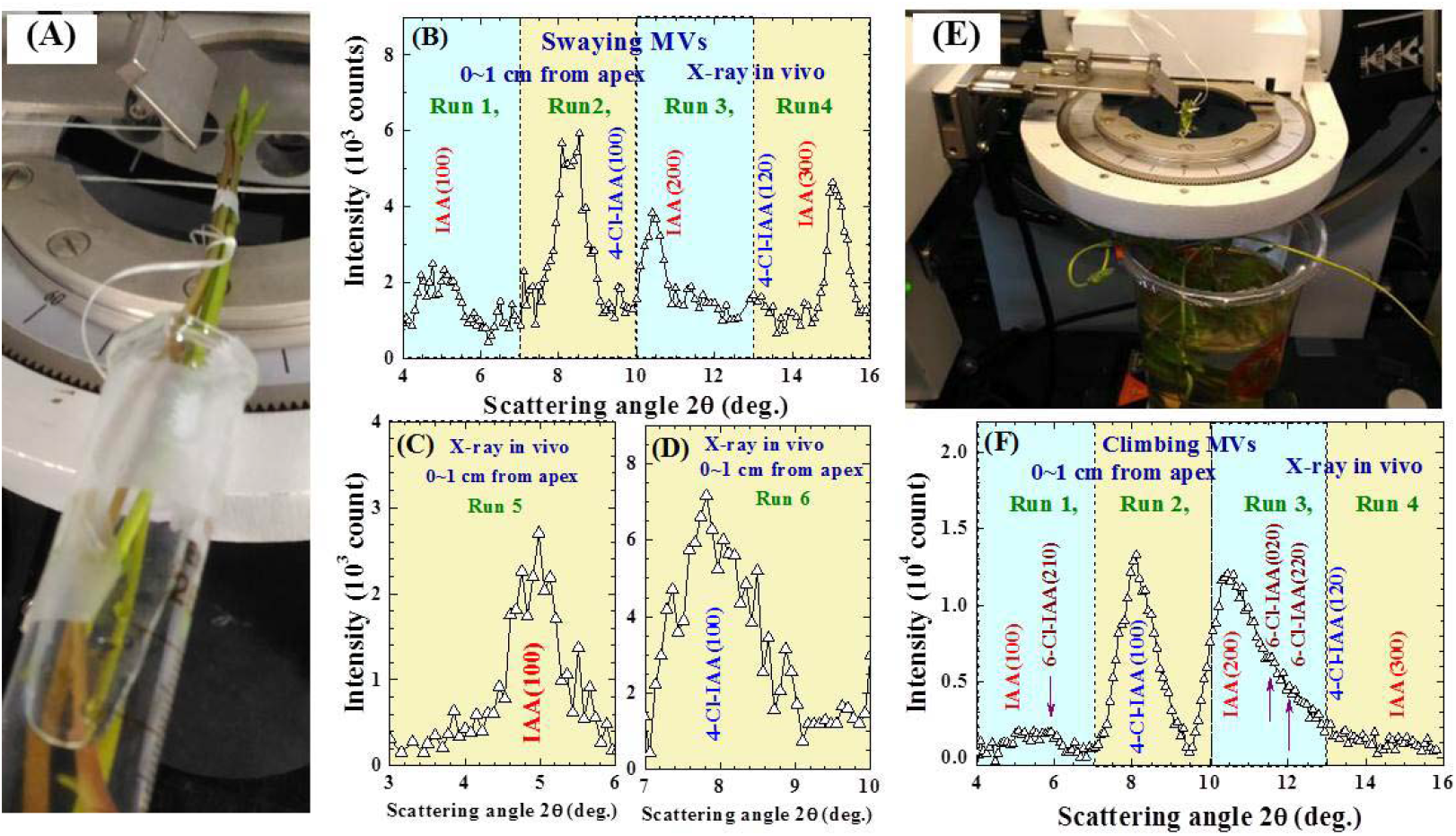
In vivo X-ray diffraction patterns. (A) Transmission geometry used in taking the X-ray diffraction patterns of a bundle of four swaying MVs. (B-D) The X-ray diffraction patterns of the twining MVs. (E) Transmission geometry used in taking the X-ray diffraction patterns of a bundle of four twining MVs. (F) The X-ray diffraction patterns of the twining MVs. Each shaded region in (B), (C), (D) and (F) indicates a 3 hour scan over a 3 degree scattering angle, followed by a beam off on MVs for 6 hrs. The Run numbers indicate the sequence used for the scans.

### Enhanced expression of 4-Cl-IAA and 6-Cl-IAA by touch stimulus

The chemical compositions of the crystallized IAA, 4-Cl-IAA and 6-Cl-IAA observed in the live MV stems were determined using the IAA extracted separately from 150 SA and 95 CA stems. Three specific portions of each CA stem were sectioned: 0-1 cm from the shoot apex in the straight (ST) part of the stems; 0-5 cm from the contact point in the curved stems bisected into contact (CT) and contact-free (CF) sides. The IAA sample extracted from a-b cm from the shoot tip of SA/CA stems was labelled SAab/CAab. Not only diffraction peaks associated to crystallized IAA but also to crystallized 4-Cl-IAA appear in the SA01 sample (crosses in Fig. 4A), while an additional series of diffraction peaks belonging to the 6-Cl-IAA also appear in the CA01 sample (open circles in Fig. 4A). The relative intensities of the 4-Cl-IAA diffraction peaks with respect to those from the IAA were significantly higher for the CA01 than for the SA01. The relative fraction of each component was obtained by employing the General Structure Analysis System (GSAS) program to quantitatively refine the XRD patterns (solid curves in Fig. 4A), following the Rietveld refinement method (Rietveld, 1969). A mass ratio of IAA:4-Cl-IAA = 96:4 is identified for SA01, and IAA:4-Cl-IAA:6-Cl-IAA = 69:28:3 for CA01, as listed in Fig. 4C.

**Fig. 4.**
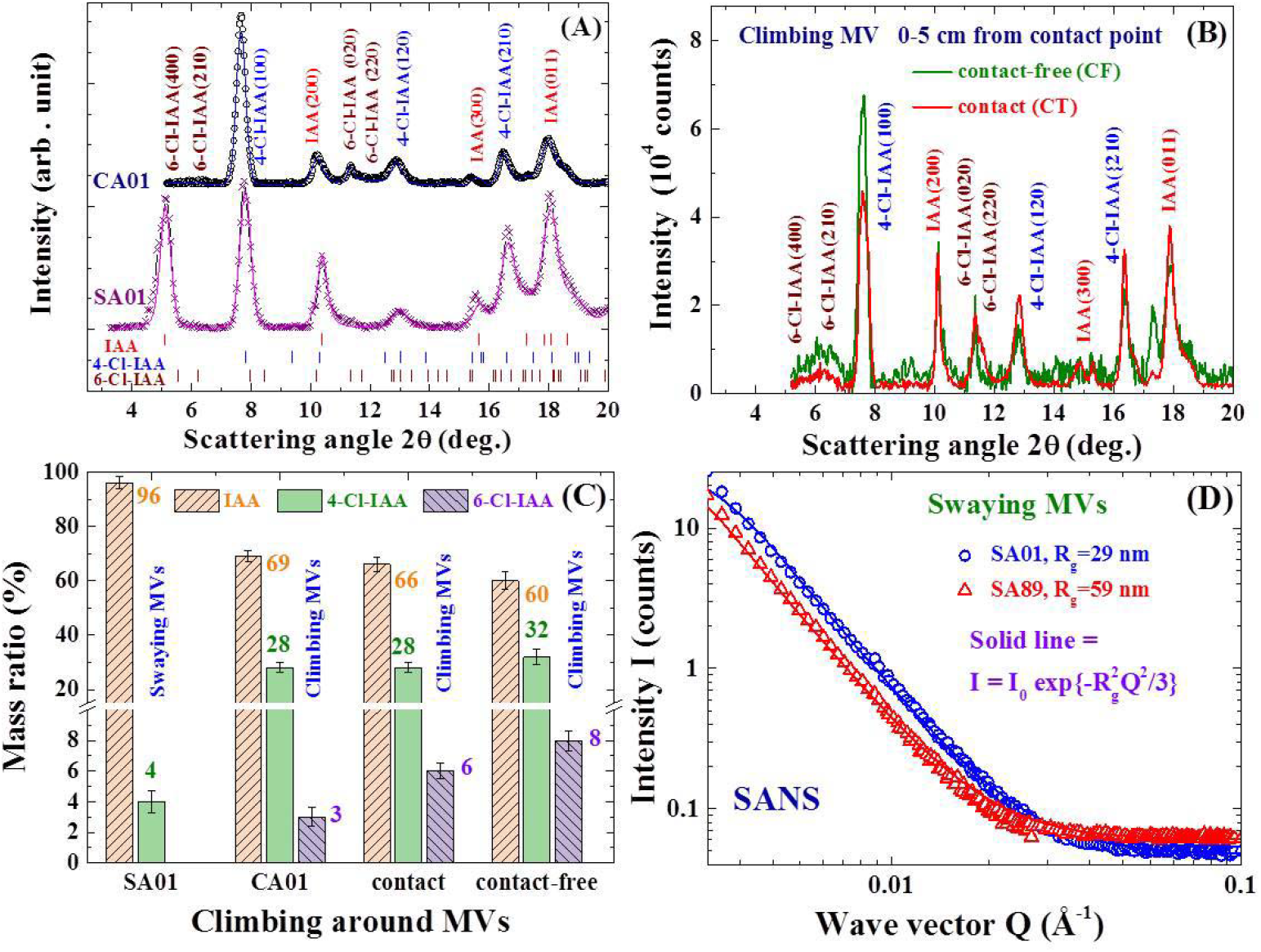
X-ray diffraction and small angle neutron scattering patterns of IAA extracted from swaying MVs. (A) Observed (crosses and open circles) and calculated (solid curves) X-ray diffraction patterns of SA01 (bottom) and CA01 (top). The vertical bars at the bottom indicate the calculated positions of the diffraction peaks. (B) X-ray diffraction patterns of IAAs extracted powder from the contact side (red curve) and the contact-free side (green curve) of 0-5 cm from the contact point of twining MVs. (C) Mass ratios of the IAA (orange bars), 4-Cl-IAA (green bars) and 6-Cl-IAA (blue bars) at various positions on the MV stems. (D) Small angle neutron scattering patterns of SA01 (open circles) and of SA89 (open triangles). The solid curves indicate the results of fits to the Guiner-Porod expression. Rg indicate the radius of gyration obtained from the fits.

Standardized against the amount of IAA, the CF side has more 4-Cl-IAA and 6-Cl-IAA than does the CT side (Fig. 4B). Refinement of the XRD patterns shows that the mass compositions of 4-Cl-IAA (green bars in Fig. 4C) and 6-Cl-IAA (blue bars in Fig. 4C) are higher on both the CF and CT sides than those in the ST part. There is more 4-Cl-IAA and 6-Cl-IAA in the curved part of the stem. The increasing amounts of 4-Cl-IAA and 6-Cl-IAA could not have been produced by the curved stem, but rather by conversion of the existing IAA as it is transported downward. The halogenation of IAA into 4-Cl-IAA and 6-Cl-IAA in the curving stem requires a Cl^-^-rich environment. The higher levels of 4-Cl-IAA and 6-Cl-IAA on the CF side reflect the appearance of a richer Cl^-^ environment than on the CT side. It has been demonstrated that monohalogenation of indole-3-acetic acid can significantly alter auxin activity. Specifically, 4-Cl-IAA is 10 times more active than IAA, and 6-Cl-IAA is 19 times more active (Böttger et al., 1978). Clearly, touch stimulus greatly enhances the expression of active 4-Cl-IAA and 6-Cl-IAA which is apparently one of the main triggers the faster growth of the MVs after touching the support.

### Elongation of auxin during transport

The sizes of IAAs in the extracted powders were determined by small angle neutron scattering (SANS), where the intensity profile *I*(Q) at the small wavevector transfer Q links directly to the radius of gyration R_g_ of the crystallized molecules in the dilute solution. The Guinier-Porod expression *I*(Q) = *I*_*0*_exp{-(R_g_ ^2^Q^2^)/3} (Guinier, 1939; Hammouda, 2010) was used to extract R_g_. The decay rate of *I*(Q) of SA89 at 20 °C (filled triangles in Fig. 4D) is obviously higher than that of SA01 (open circles in Fig. 4D), showing that the IAA/4-Cl-IAA in the stem 8-9 cm below the shoot apex had crystallized to larger sizes than 0-1 cm below the apex. Fits of the *I*(Q) to the Guinier-Porod expression (solid curves in Fig. 4D) give *R*_g_ = 29(2) nm for the IAA/4-Cl-IAA in the area 0-1 cm below the shoot apex and 59(3) nm in the area 8-9 cm below the apex. Note that *R*_g_ indicates the periodically aggregated size of IAA/4-Cl-IAA. The *R*_g_ of the crystallized IAA/4-Cl-IAA becomes larger in the lower portion of the stem toward the root. The natural biosynthesis of IAA/4-Cl-IAA at the shoot apex is likely to occur in molecular form. Upon transport downward through the cells, periodic aggregation of the molecular IAA/4-Cl-IAA into crystallized form occurs, with the size of the aggregated becoming larger in the lower portion toward the root of the stem.

### Bio-environment for differential growth

Taking the advantage of the high sensitivity of neutron to hydrogen, in vivo neutron tomography of a bundle of naturally climbing MVs (Fig. 5A) was carried out to examine the hydrogen distribution in the living MVs. Figures 5B and 5C display representative slices from the tomographic images within which contact between MVs appeared. The colors in the stems represent the neutron intensity, with the uncolored white areas represent the base color for minimum intensity. Obvious differences in the neutron intensity, hence the hydrogen content, between the CT side (in grey) and the CF side (in yellow) are clearly reviewed. A well-shaped intensity distribution of the hydrogen content across the stem is seen, with the well extending over ~50% of the stem (Fig. 5D). No obvious difference in the hydrogen content across the stem is seen for the stems that are isolated from each other (Figs. 5E, 5F, 5G). Clearly, contact between MVs triggers a redistribution of the hydrogen in the stem, with a portion of the hydrogen on the CT side shifting to the CF side. The appearance of more hydrogen on the CF side is also reflected by the higher pH value obtained for the CF side than for the CT side (Fig. 6A), with the concentration of H^+^ on the CF side being (10^6.77-6.74^)-1 = 9% higher. No obvious difference was detected between the H^+^ concentrations in the area 0-1 cm down from the apex of the SA and CA MVs, with the pH values for these two parts being essentially the same (Fig. 6A). It is known that H^+^ will facilitate transportation of IAA into cells.

**Fig. 5.**
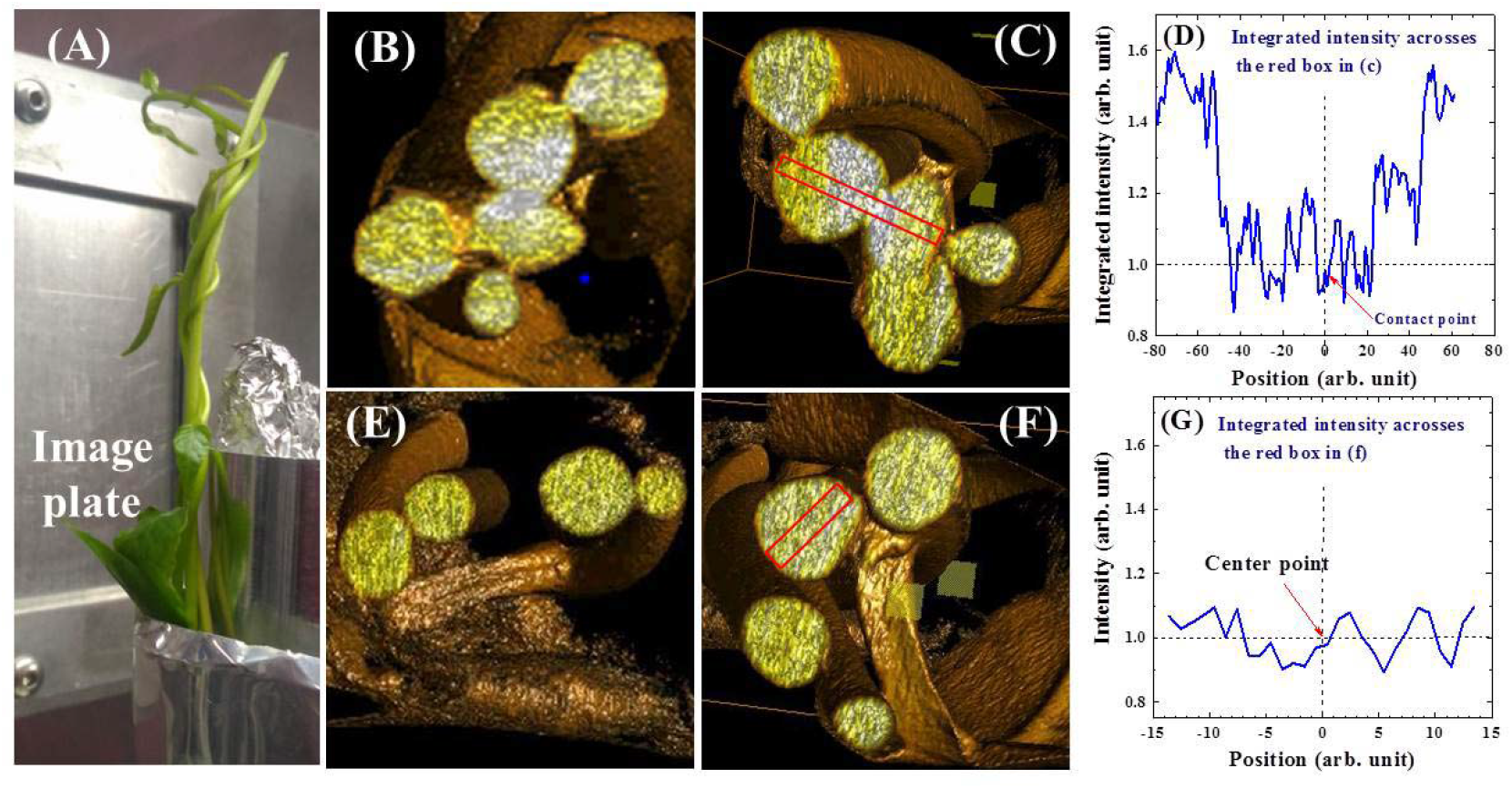
Neutron tomography of live MVs. (A) Bundle of MVs used for taking neutron tomography measurements. (B)-(C) Images from the tomography reveal contact between MVs. (D) Integrated intensities across the red box marked in (C), revealing a well-shaped intensity distribution. (E)-(F) Images from the tomography revealing isolated MVs. (G) Integrated intensities across the red box marked in (F).

**Fig. 6.**
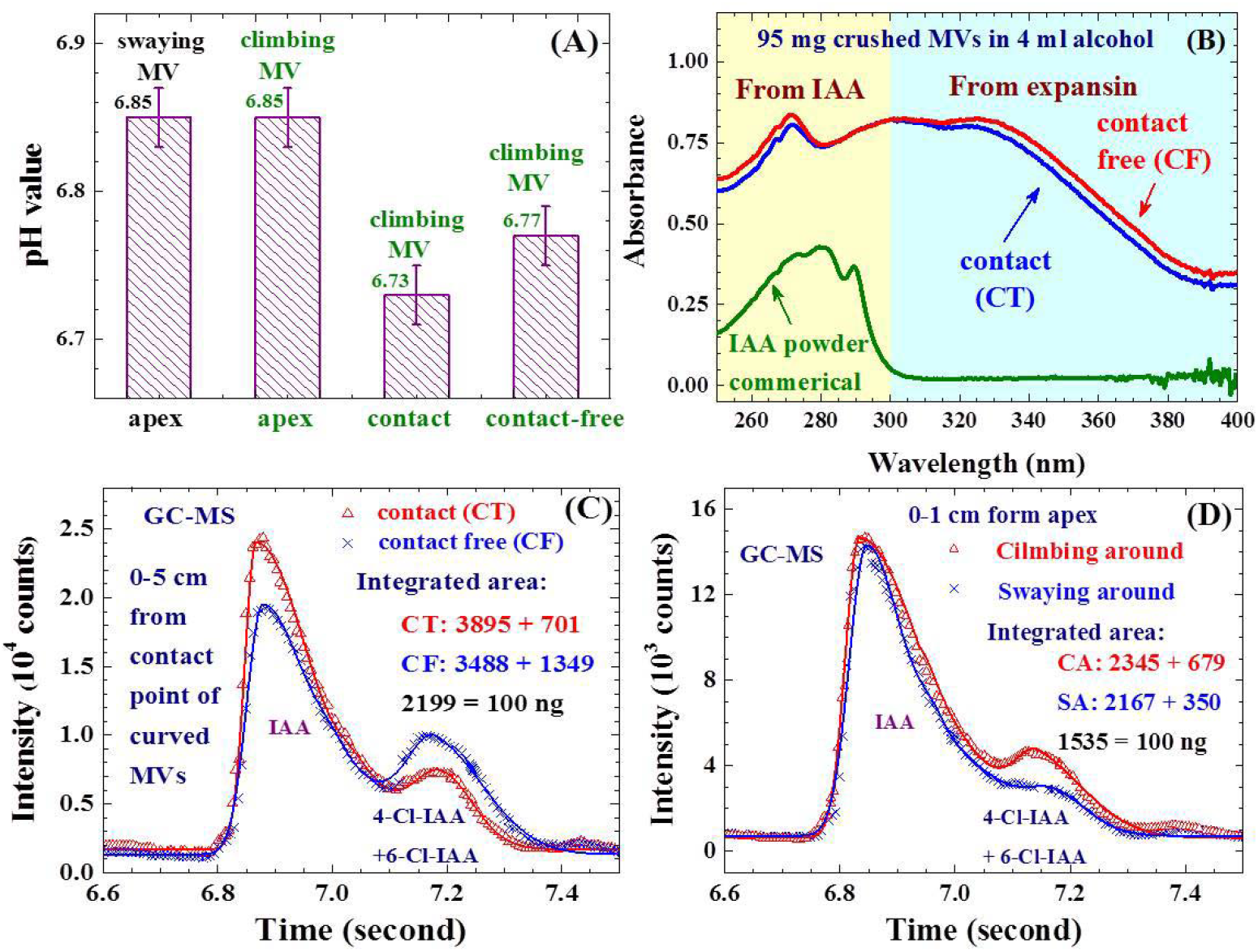
Chemical environment in contact-free and contact sides of curved MVs. (A) pH values at various positions in the swaying around and twining MVs. (B) Direct comparisons between absorption spectra of commercial IAA (green curve), contact-free side (red curve) and contact side (blue curve) of freshly crashed curved MVs. The green shaded region marks the absorption from IAA, and the blue shaded region marks the absorption from expansin. (C) GC-MS spectra of the IAAs extracted from the contact-free side (crosses) and contact side (open triangles) of 0-5 cm from the contact point on curved MVs. (D) GC-MS spectra of the IAAs extracted from the area 0-1 cm from the shoot apex of twining (open triangles) and swaying (crosses) MVs.

Two components appear in the absorption spectra of crushed MV stems mixed in alcohol (Fig. 6B). The component covering 250-300 nm with a peak at ~270 nm reflects the absorption by the IAA, whereas the one covering 300-400 nm with peaks at ~310 and 330 nm is linked to the absorption by expansin (Tabuchi et al., 2011). More IAA and expansin appear on the CF side than on the CT side (Fig. 6B). The IAA and expansin on the CF side are respectively 4% and 6% higher than those on the CT side, when estimated using the integrated absorbance of the two components. Expansins are plant cell-wall loosening proteins that facilitate the transportation of auxin into cells for cell elongation (Tabuchi et al., 2011).

The amount of IAA in the stems was measured using gas chromatography–mass spectrometry (GC-MS) performed on the HPLC powders. In addition to the peak observed at the time channel of t ~ 6.84 sec for IAA, a separated peak at t ~ 7.14 sec was also clearly evident in the GC-MS spectra of the powders extracted from the apex region (0-1 cm from the apex) of the CT (open triangles in Fig. 6C)) and CF stems (crosses in Fig. 6C). The peak at t ~7.14 sec is linked to 4-Cl-IAA and 6-Cl-IAA. As they have the same molecular weight but are 17% heavier than IAA to appear at a slightly later time in the spectrum. An examination of the two spectra shown in Fig. 6C reveal the appearance of visibly larger amount of IAA together with a significantly larger amount of 4-Cl-IAA+6-Cl-IAA in the CA stems than in the SA stems. The absolute value of the IAA can then be obtained by comparing the integrated intensity of the associated peak with that obtained from 100 ng of standard IAA (from Sigma-Aldrich). We found 153 ng of IAA and 44 ng of 4-Cl-IAA+6-Cl-IAA in each gram of dried stems in the sample 0-1 cm from the apex of the CA MVs, and 141 ng of IAA and 23 ng of 4-Cl-IAA+6-Cl-IAA for the SA MVs (Table 1). Touch stimulus triggered the apex to produce 9% more IAA together with 91% more 4-Cl-IAA+6-Cl-IAA. In addition, the GC-MS spectra revealed a significantly lower amount of IAA accompanied by a significantly larger amount of 4-Cl-IAA+6-Cl-IAA on the CF side than on the CT side (Fig. 6D). Comparing the integrated intensity to the standard IAA powder give 159 ng of IAA together with 61 ng 4-Cl-IAA+6-Cl-IAA that appear in each gram of dried stems from the CF side, and 177 ng of IAA and 32 ng 4-Cl-IAA+6-Cl-IAA from the CT side (Table 1). About 220 ng of IAA+4-Cl-IAA+6-Cl-IAA appear on the CF side and 209 ng on the CT side. Physical contact drives the apex to produce significantly more IAA (~9% more) and 4-Cl-IAA+6-Cl-IAA (~90% more), but transport more 4-Cl-IAA+6-Cl-IAA to the CF side where more H^+^ ions and expansins have accumulated. This in turn facilitates the entry of IAA+4-Cl-IAA+6-Cl-IAA into the cell, which gives rise to the differential growth allowing the stem to curve around the support.

**Table 1.** Specific masses of IAA and 4-Cl-IAA+6-Cl-IAA per gram of dried stem at various positions along the MV stems.

|  | IAA (ng) | 4-Cl-IAA+6-Cl-IAA (ng) | Total (ng) |
| --- | --- | --- | --- |
|  | Per gram of dried stem |  |  |
| Swaying MVs 0-1 cm from apex | 141 | 23 | 164 |
| Climbing MVs 0-1 cm from apex | 153 | 44 | 197 |
| Contact side 0-5 cm from contact point | 177 | 32 | 209 |
| Contact-free side 0-5 cm from contact point | 159 | 61 | 226 |

## Discussion

Madeira vine (MV) is an evergreen vine with a worldwide distribution. MV sways around counterclockwise at a frequency of typically 5 hrs per turn. The shoot apex produces IAA and 4-Cl-IAA supporting a growth rate of 2.4 mm/day while swaying around, with no considerable difference in swaying frequency or in growth speed between daytime and nighttime. Upon being transported downward: (1) the IAA and 4-Cl-IAA molecules link to form a crystallized arrangement, reaching 29 nm in size 0-1 cm below the shoot apex and 59 nm in the area 8-9 cm below; (2) a portion of the IAA chlorinated to form 4-Cl-IAA, with a mass proportion of 4-Cl-IAA:IAA increasing from 1:19 in 0-1 cm below the shoot apex to 3:1 in 8-9 cm below.

MV is a fast climber upon making physical contact with a support. It takes ~3 days after contacted with a support for MV to reach the fastest growth of 9 mm/12 hrs in the daytime (7 am to 7 pm) and 35 mm/12 hrs in the nighttime (7 pm to 7 am). The 4-fold faster growth speed during the nighttime demonstrates that cell elongation and division slow down during the daytime. It takes ~1 day for MV to grow an additional ~20 mm after detachment from the support to resume the swaying around growth pattern. The apex lose the stimulation of physical contact once it is 20 mm away from the contact point. Physical contact stimulates (1) the apex to produce 6-Cl-IAA in addition to 10% more IAA and double the expression of 4-Cl-IAA, with a mass proportion of 6-Cl-IAA:4-Cl-IAA:Cl-IAA = 3:28:69; (2) the transport of more 4-Cl-IAA and 6-Cl-IAA toward the contact-free side; (3) a higher chlorination rate of IAA to form 4-Cl-IAA and 6-Cl-IAA, with a mass proportion of 6-Cl-IAA:4-Cl-IAA:Cl-IAA = 6:28:66 on the contact side and 8:32:60 on the contact-free side; (4) the diffusion of 9% more H^+^ and a 6% more expansin from the contact side toward the contact-free side, presumably trigged by the higher mechanical pressure created by physical contact.

Thigmotropism is initiated by the physical touch of the stem to a support. The touch transfers the energy used in swaying around to produce faster growth. Climbing or twining around a support is the natural way for plant stems to remain in contact but this needs differential growth. In the Madeira vine, faster growth is linked to the enhanced expression of 4-Cl-IAA and 6-Cl-IAA, and the differential growth is linked to the transport of more 4-Cl-IAA and 6-Cl-IAA together with shifting of the H^+^ and expansin from the contact side toward the contact-free side.

## Author contributions

W.H.L. and M.H.M. designed the study; J.-D.C., C.-T.C. and Y.-H.T. perform HPLC extractions; M.H.M., E.B., H.-H.L., C.-I.L. and N.-J.C. performed the measurements; C.M.W., M.H.M. and W.H.L. performed the neutron measurements; M.H.M., E.B., H.-H.L. and W.H.L. analyze the data; all of the authors discussed the results; M.H.M. and W.H.L. wrote the manuscript.

## References

Ahmad A, Anderson AS, Engvild K. 1987. Rooting, growth and ethylene evolution of pea cuttings in response to chloroindole auxins. Physiologia Plantarum 69, 137–140.

Bennett T, Hines G, van Rongen M, Waldie T, Sawchuk MG, Scarpella E, Ljung K, Leyser O. 2016. Connective auxin transport in the shoot facilitates communication between shoot apices. PLoS Biology 14, e1002446.

Bennett T, Sieberer T, Willett B, Booker J, Luschnig C, Leyser O. 2006. The Arabidopsis MAX pathway controls shoot branching by regulating auxin transport. Current Biology 16, 553–563.

Bopp M, Weber I. 1981. Hormonal regulation of the leaf blade movement of Drosera capensis. Physiologia Plantarum 53, 491–496.

Böttger M, Engvild KC, Soll H. 1978. Growth of Avena coleoptiles and pH drop of protoplast suspensions induced by chlorinated indoleacetic acids. Planta 140, 89–92.

Bowling AJ, Vaughn KC. 2009. Gelatinous fibers are widespread in coiling tendrils and twining vines. American Journal of Botany 96, 719–727.

Braam J. 2005. In touch: plant responses to mechanical stimuli. New Phytologist 165, 373–389.

Burris JN, Lenaghan SC, Stewart CN. 2018. Climbing plants: attachment adaptations and bioinspired innovations. Plant Cell Reports 37, 565–574.

Darwin C. 1881. The power of movement in plants. New York: D. Appleton and Company.

Esmon CA, Pedmale UV, Liscum E. 2005. Plant tropisms: providing the power of movement to a sessile organism. International Journal of Developmental Biology 49, 665–674.

Jaffe MJ, Galston AW. 1968. The physiology of tendrils. Annual Review of Plant Physiology 19, 417–434.

Jaffe MJ, Leopold AC, Staples RC. 2002. Thigmo responses in plants and fungi. American Journal of Botany 89, 375–382.

Gälweiler L, Guan C, Müller A, Wisman E, Mendgen K, Yephremov A, Palme K. 1998. Regulation of polar auxin transport by AtPIN1 in Arabidopsis vascular tissue. Science 282, 2226–2230.

Gianoli E. 2015. The behavioural ecology of climbing plants. AoB Plants 7.

Guinier A. 1939. La diffraction des rayons X aux très petits angles: application à l’étude de phénomènes ultramicroscopiques. Annales de physique 11, 161–237.

Guerra S, Peressotti A, Peressotti F, et al. 2019. Flexible control of movement in plants. Scientific Reports 9, 16570.

Hammouda B. 2010. A new Guinier-Porod model. Journal of Applied Crystallography 43, 716–719.

Iino M. 2006. Toward understanding the ecological functions of tropisms: interactions among and effects of light on tropisms. Current Opinion in Plant Biology 9, 89–93.

Karcz W, Burdach Z. 2002. A comparison of the effects of IAA and 4-Cl-IAA on growth, proton secretion and membrane potential in maize coleoptile segments. Journal of Experimental Botany 53, 1089–1098.

Korasick DA, Enders TA, Strader LC. 2013. Auxin biosynthesis and storage forms. Journal of Experimental Botany 64, 2541–2555.

Nigović B, Kojić-Prodić B, Antolić S, Tomić S, Puntarec VITIMIR, Cohen JD. 1996. Structural studies on monohalogenated derivatives of the phytohormone indole-3-acetic acid (auxin). Acta Crystallographica Section B: Structural Science 52, 332–343.

Perrot-Rechenmann C. 2010. Cellular responses to auxin: division versus expansion. Cold Spring Harbor Perspectives in Biology 2, a001446.

Pless T, Böttger M, Hedden P, Graebe J. 1984. Occurrence of 4-Cl-indoleacetic acid in broad beans and correlation of its levels with seed development. Plant Physiology 74, 320–323.

Rietveld HM. 1969. A profile refinement method for nuclear and magnetic structures. Journal of Applied Crystallography 2, 65–71.

Rowe N, Speck T. 2005. Plant growth forms: an ecological and evolutionary perspective. New Phytologist 166, 61–72.

Scorza LC, Dornelas MC. 2015. Exploring the role of auxin in the androgynophore movement in Passiflora. Genetics and Molecular Biology 38, 301–307.

Smyth DR. 2016. Helical growth in plant organs: mechanisms and significance. Development 143, 3272–3282.

Steinitz B, Hagiladi A. 1987. Thigmomorphogenesis in Climbing Epipremnum aureum, Monsters obliqua expilata and Philodendron scandens (Araceae). Journal of Plant Physiology 128, 461–466.

Stolarz S. 2009. Circumnutation as a visible plant action and reaction. Plant Signaling & Behavior 4, 380–387.

Tabuchi A, Li LC, Cosgrove DJ. 2011. Matrix solubilization and cell wall weakening by β-expansin (group-1 allergen) from maize pollen. Plant Journal 68, 546–559.

Tivendale ND, Davidson SE, Davies NW, Smith JA, Dalmais M, Bendahmane AI, Quittenden LJ, Sutton L, Bala RK, Le Signor C, Thompson R, Horne J, Reid JB, Ross JJ. 2012. Biosynthesis of the halogenated suxin, 4-chloroindole-3-acetic acid. Plant Physiology 159, 1055–1063.

Vieten A, Sauer M, Brewer PB, Friml J. 2007. Molecular and cellular aspects of auxin-transport-mediated development. Trends in Plant Science 12, 160–168.

Zhang Y, Wu J, Ma Y, Chen H, Chen Y, Lu B, Feng X. 2018. A finite deformation theory for the climbing habits and attachment of twining plants. Journal of Mechanics and Physics of Solids 116, 171–184.

Živanović BD, Ullrich KK, Steffens B, Spasić SZ, Galland P. 2018. The effect of auxin (indole-3-acetic acid) on the growth rate and tropism of the sporangiophore of Phycomyces blakesleeanus and identification of auxin-related genes. Protoplasma 255, 1331–1347.

